# A hidden Markov model approach to characterizing the photo-switching behavior of fluorophores

**DOI:** 10.1101/223875

**Authors:** Lekha Patel, Nils Gustafsson, Yu Lin, Raimund Ober, Ricardo Henriques, Edward Cohen

**Author notes:** Corresponding author. Dr Edward Cohen, Department of Mathematics, Imperial College London, South Kensington Campus, London SW7 2AZ.

## Abstract

Fluorescing molecules (fluorophores) that stochastically switch between photon-emitting and dark states underpin some of the most celebrated advancements in super-resolution microscopy. While this stochastic behavior has been heavily exploited, full characterization of the underlying models can potentially drive forward further imaging methodologies. Under the assumption that fluorophores move between fluorescing and dark states as continuous time Markov processes, the goal is to use a sequence of images to select a model and estimate the transition rates. We use a hidden Markov model to relate the observed discrete time signal to the hidden continuous time process. With imaging involving several repeat exposures of the fluorophore, we show the observed signal depends on both the current and past states of the hidden process, producing emission probabilities that depend on the transition rate parameters to be estimated. To tackle this unusual coupling of the transition and emission probabilities, we conceive transmission (transition-emission) matrices that capture all dependencies of the model. We provide a scheme of computing these matrices and adapt the forward-backward algorithm to compute a likelihood which is readily optimized to provide rate estimates. When confronted with several model proposals, combining this procedure with the Bayesian Information Criterion provides accurate model selection.

## 1 INTRODUCTION

Fluorescence microscopy is a collection of techniques that utilize the photon emitting properties of fluorescing molecules, called *fluorophores*, to perform optical imaging, particularly in cell biology and biomedical applications. Recent years have seen the advent of a number of super-resolution microscopy techniques that have bypassed the classical resolution limits of fluorescence microscopy (Huang et al., 2009). Specifically, single molecule localization microscopy (SMLM) approaches, such as photoactivated localization microscopy (PALM) (Betzig et al., 2006; Hess et al., 2006) and stochastic optical reconstruction microscopy (STORM) (Rust et al., 2006; Heilemann et al., 2008), rely on the ability exhibited by some fluorophores to *photoswitch* stochastically between a photon emitting On state and non-emitting dark states (Van de Linde and Sauer, 2014; Ha and Tinnefeld, 2012). A specimen decorated with a spatially dense number of photon emitting fluorophores prevents accurate identification of individual fluorophores and resolution of structures smaller than the diffraction limit – see Figure 1 (a). Using a fluorophore with stochastic photo-switching properties can provide an imaging environment where the majority of fluorophores are in a dark state, while a sparse number have stochastically switched into a transient photon emitting On state. This results in the visible fluorophores being sparse and well separated in space; with the use of a high-performance camera the individual fluorophores in the On state can be identified and localized with nanometer scale precision by fitting point spread functions (Sage et al., 2015; Ober et al., 2015) – see Figure 1 (b). Through the acquisition across time of a large sequence of images (typically thousands) – see Figure 1 (a) – many more photo-switching fluorophores can be isolated in time and precisely localized in space. When aggregated and plotted, these localizations provide an accurate and detailed map of fluorophore positions giving rise to a super-resolved image.

**Figure 1:**
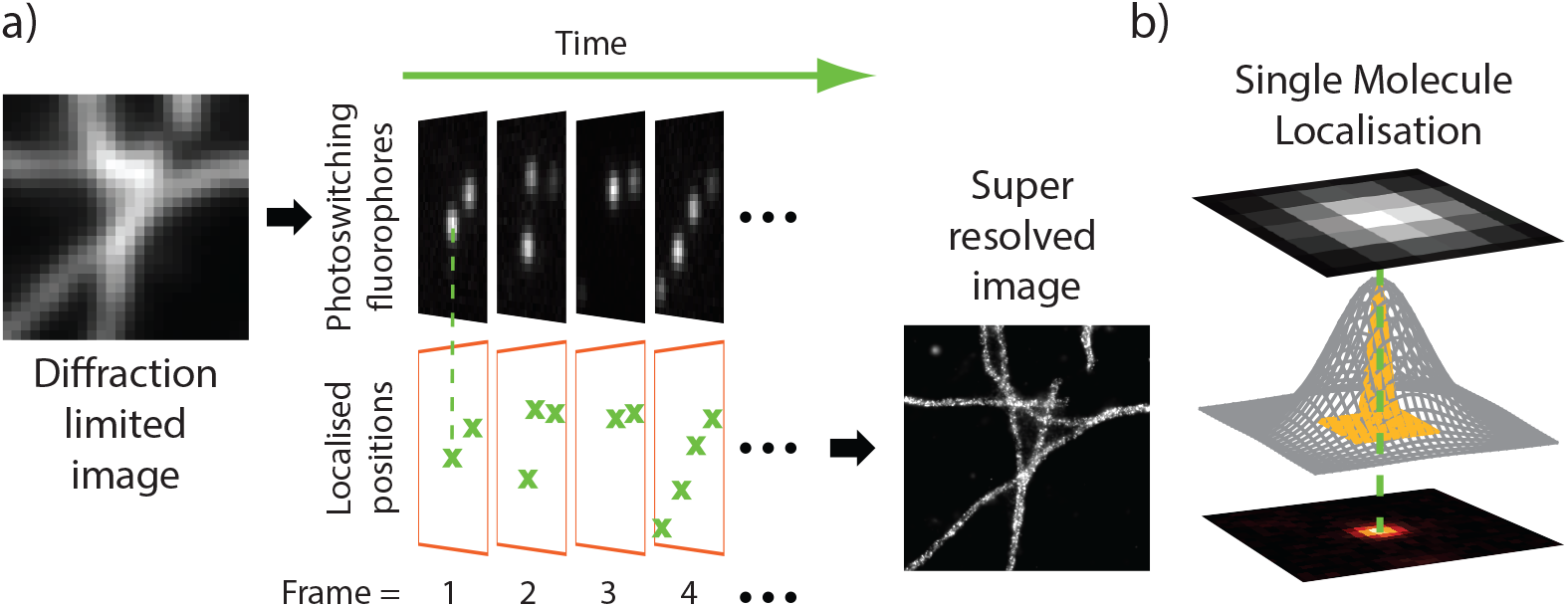
(a) Illustration of the SMLM imaging process. When all fluorophores simultaneously stay in a photon emitting On state, diffraction renders structures unresolvable. Stochastically photo-switching fluorophores imaged over time across several frames give rise to a sequence of sparsely populated images where each fluorophore can be isolated and localized with high precision. Aggregating these frames gives rise to a super-resolved image. Data from Sage et al. (2015) (b) Isolated fluorophores are localized by fitting the point spread function (PSF) to the diffraction limited spot.

While lateral resolutions of 10 – 30 nanometers (nm) are possible in biological samples using SMLM, the resolution and image quality is strongly dependent on the photo-switching properties of the fluorophore used. This dependence arises primarily because longer On states provide a greater number of photons being recorded by the camera, which in turn leads to greater precision in localizing fluorophores (Ober et al., 2004; Ram et al., 2012; Thompson et al., 2002; Rieger and Stallinga, 2014). However, the increased random occurrence of fluorophores simultaneously occupying the On state within a diffraction limited spot can lead to significant imprecision, missed events and unwanted artifacts (Van de Linde et al., 2010; Nieuwenhuizen et al., 2015). Thus, a careful choice of fluorophore and the environment used to promote photo-switching – controlled by the buffer solution and illumination intensity – must be made for the intended application. This is particularly important in live-cell applications when considerations must be made for temporal resolution and reduced laser intensities.

In order to inform the choice of fluorophore with its environment, and aid the development of novel fluorophores, accurate characterization of the photo-kinetic model of the fluorophore, together with estimation of photo-switching rates (the rate at which fluorophores transition between On and dark states) is required (Dempsey et al., 2011; Lehmann et al., 2016). Further to this, accurate knowledge of the photo-switching characteristics could be employed to maximize resolutions achieved using advanced analytical methods, for example, 3B analysis (Cox et al., 2011) and DeconSTORM (Mukamel et al., 2012) and improve the performance of molecular counting techniques (Rollins et al., 2014; Lee et al., 2012).

A number of attempts have been made to model the kinetic schemes of fluorophore photoswitching and estimate the corresponding photo-switching rates. These kinetic schemes, as is common across single molecule biophysics, are characterized by Markovian transitions between a finite set of discrete states and are therefore ideally suited to being modeled as continuous time Markov processes. In Figure 2 are four models for photo-switching fluorophores. The first, 2a, depicts a typical kinetic model, accompanied by the state-space diagram we will adopt in this paper. In this model there is a photon emitting On state 1 (involving rapid transitions between excited state S_1_ and ground state S_0_ via the absorption and emission of a photon), two temporary dark states 0 and 0_1_ (the triplet state, T_1_ and the redox states, F^+^ and F^−^) and an absorption 2 (BF/BT_0_/BS_0_) which in this application is known as the *photobleached* state. Then in Figures 2b-2d are three further common state space models. Figure 2b portrays a photo-switching model with a simple two state {On(1) Dark(0)} structure. Models of this type are suitable for super-resolution methods including point accumulation for imaging in nanoscale topography (PAINT) and DNA-PAINT (Sharonov and Hochstrasser, 2006; Jungmann et al., 2010). Figure 2c depicts a model that incorporates an absorbing state 2. This form of photo-switching followed by absorption describes a first approximation to the behavior that occurs spontaneously in a number of organic fluorophores and post-activation of photoactivatable proteins (Van de Linde and Sauer, 2014; Ha and Tinnefeld, 2012; Vogelsang et al., 2010). Figure 2d considers a model in which three distinct dark states are hypothesized which in some cases is a necessary extension to model (c), for instance when very rapid imaging is used (Lin et al., 2015).

**Figure 2:**
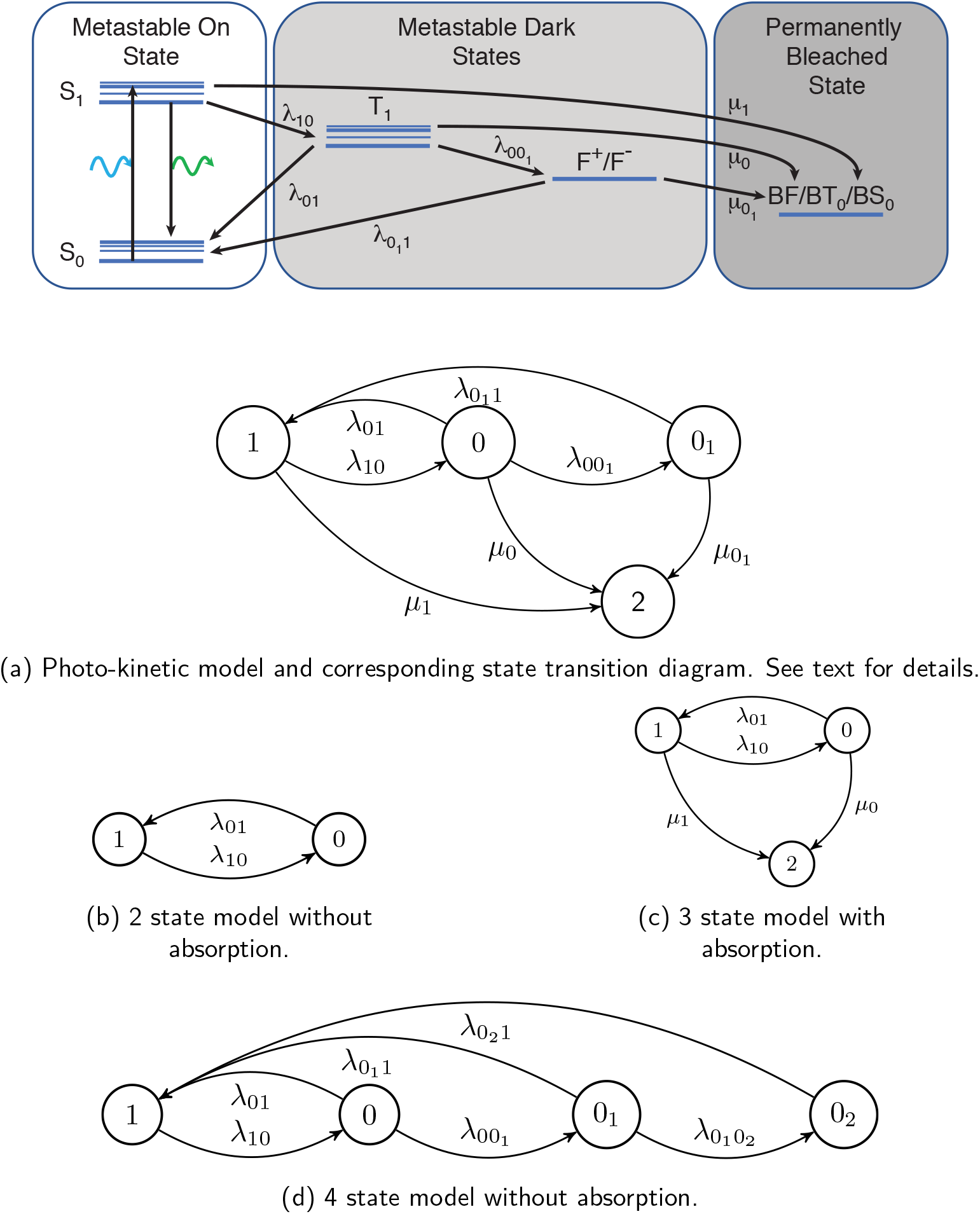
Common models used to describe the continuous time photo-switching dynamics of a fluorophore with homogeneous transition rates. See text for details.

The challenge comes in selecting the correct model and estimating the transition rates of the continuous time Markov process 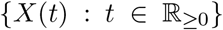 from an observed discrete-time random process 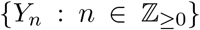. Here, 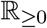 and 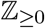 denote the non-negative reals and integers, respectively. Typically, {*Y_n_*} is derived from a sequence of images (frames) with *Y_n_* corresponding to the *observed* state of the molecule in the *n*th frame. This is formed by an exposure of the continuous time process {*X*(*t*)} over the time-interval [*n*Δ, (*n* + 1)Δ), where Δ is the frame length. Process {*Y_n_*} can either be a sequence of photon fluxes associated with that molecule for each frame (Figure 3a), or a simple sequence of 1s and 0s indicating if the molecule was detected in the frame or not (Figure 3b). In all cases, the observations are subject to the effects of noise and instrument limitations. Essential to the subsequent analysis, therefore, is the ability to account for missed state transition events due to noise and the temporal resolution of the data acquisition, as well as the detection threshold used to determine the state of the system (Figure 3c). Similar problems occur in other areas of biophysics where estimating transition rates of an underlying continuous time Markov process must be inferred from an observed discrete time signal. In particular, ion-channels have formed the focus of much work (Colquhoun and Hawkes, 1981; Qin and Li, 2004; Rief et al., 2000), including methods that attempt to account for missed events (Qin et al., 1996; Colquhoun et al., 1996; Hawkes et al., 1990, 1992; Epstein et al., 2016). However, the mechanism by which the observed signal is obtained and processed from the raw signal is fundamentally different to that of fluorescence microscopy imaging.

**Figure 3:**
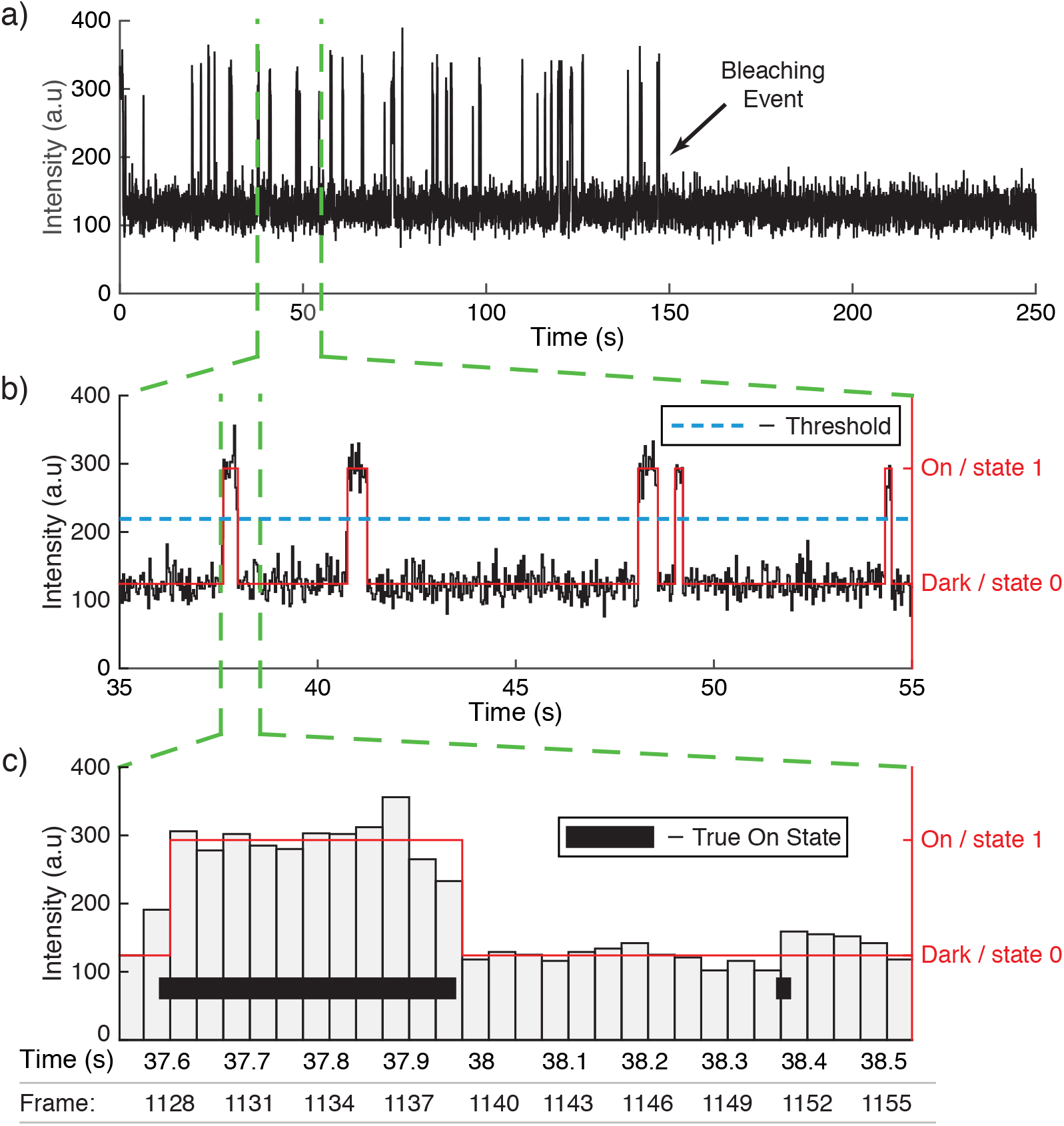
(a) A simulated intensity signal of a fluorophore across time. Each measurement corresponds to the intensity in a frame. 7500 frames were recorded over 250 seconds at a rate of 30 frames per second. (b) Close up of the signal over the time window of 35s to 55s. In red is the observed signal {*Y_n_*} indicating if the fluorophore was detected in a particular frame. (c) A further close up of the signal showing intensity read-outs for independent frames. The true, hidden photon emitting On state of the molecule is also indicated, demonstrating how sub-frame length photon emitting events can be missed due to noise or the temporal resolution of the data acquisition.

Up until now, methods for estimating photo-switching transition rates in fluorescence microscopy are limited. The method in Lin et al. (2015) involves defining {*Y_n_*} to be the sequence of 1s and 0s and extracting the dwell times. These are the set of time lengths that {*Y_n_*} is seen to sit in the On state and the set of time lengths it is seen to sit in a dark state. Assuming these dwell times to be exponentially distributed (or a mixture of exponential distributions in the case of multiple dark states), maximum likelihood estimates of the transition times are then computed. This method, which will be referred to here as *exponential fitting*, has two flaws. Firstly, it does not correctly account for the effect the imaging procedure has on the stochastic structure of the discrete time process. Secondly, it does not allow for the absorbing (photobleached) state, which must be identified and accounted for by truncation of the data to the last observed On state. This is especially troublesome as, to an observer, it is indistinguishable from a temporary dark state. This method therefore results in the absence of estimates for the absorption rate and can lead to significantly biased estimates of the transition rates between On and dark states.

Hidden Markov models (HMMs) are widely used across a range of scientific and engineering disciplines to relate a sequence of observations, called emissions, to the states of an unobserved (hidden) Markov process that one wishes to perform inference on. Their use is particularly prevalent in image processing where the observations are a sequence of images in time and it is commonly assumed that each image is dependent only on the state of the hidden process at the time at which it is observed. Such an approach has been proposed for this problem in Greenfeld et al. (2015), where the hidden process is a discretized version of {*X*(*t*)}. Here, they let {*Y_n_*} be the sequence of photon-fluxes such that it is a standard (first-order) HMM with Poisson emissions. They then implement the Baum-Welch algorithm (Baum and Petrie, 1966; Baum and Eagon, 1967; Baum and Sell, 1968; Baum et al., 1970) to estimate the transition probabilities of the discretized process and use an approximation to obtain the transition rates of the continuous time process {*X*(*t*)}. In doing so, they acknowledge that missed events will heavily bias rate estimates. Furthermore, their model is also unable to deal with the absorbing state.

In this paper we provide two important contributions. Firstly, in Section 2, by considering a general model for {*X*(*t*)} that includes multiple dark states and an absorbing state, we rigorously formulate the discrete time stochastic process {*Y_n_*} that indicates whether a molecule is detected in each frame. A crucial part of this formulation is recognizing that a image is not formed from an instantaneous sampling of the true state, as is usually assumed in image processing, but is instead formed by exposing a camera sensor over a time interval of length Δ. That is to say, *Y_n_* is not dependent on just *X*(*n*Δ), but instead on the integral (i.e. all values) of {*X*(*t*)} over the interval [*n*Δ, (*n* + 1)Δ). Taking consideration of noise and instrument sensitivity, we fully account for missed events and give important results on the stochastic structure of {*Y_n_*}, including showing it is non-Markovian.

The second contribution of this paper is to propose novel methodology for estimating the state transition rates of {*X*(*t*)} under this correct treatment of the imaging procedure. In Section 3, we develop an HMM for {*Y_n_*} where we first implement a time discretization scheme on the hidden Markov process {*X*(*t*)}. Crucially, as discussed above, correct understanding of the imaging procedure dictates two key properties. Firstly, *Y_n_* depends on both the current (end of frame) and previous (beginning of frame) hidden states, *X*((*n* + 1)Δ) and *X*(*n*Δ), respectively. Secondly, this HMM possesses emission probabilities that are dependent on the static parameters of the hidden process state transitions that we ultimately wish to estimate. This *coupled* behavior renders traditional expectation maximization (EM)-type methods (e.g. Baum et al. (1970)) of parameter estimation inappropriate. We therefore make the novel step of introducing what we call *transmission* (transition-emission) matrices that incorporate this coupling between transition and emission probabilities by capturing all the dependencies in the model. For a given photo-switching kinetic model, we provide both a scheme for computing these matrices and an adaptation of the forward-backward algorithm to compute the likelihood of observations. Through numerical optimization we are able to compute maximum likelihood estimates of the transition rate parameters for the continuous time process {*X*(*t*)} that we wish to draw inference on. A bootstrapping scheme is also presented for computing confidence intervals. In the case of an unknown kinetic model, we propose the use of the Bayesian information criterion (BIC) for selecting the best suited model from a set of proposals, thus also providing a powerful tool for chemists wishing to infer the number of quantum states a particular fluorophore can exist in.

In Section 4, we provide a simulation study that compares this new estimation scheme to the exponential fitting method on a range of photo-switching models, demonstrating significant improvements in both the bias and the variance of our rate estimates. We further show the BIC performs accurate model selection when presented with a range of model proposals. In Section 5, the estimation scheme presented in this paper is applied to the Alexa Fluor 647 data originally analysed by the exponential fitting method in Lin et al. (2015), consistently selecting the hypothesised three temporary off-state model (Figure 2d) and revealing clear dependence between laser intensity and key transition rate parameters. In the accompanying supplementary material, as well as key mathematical details, we include an extensive simulations section where we report a significant improvement on rate estimates across a range of models and relevant experimental conditions.

## 2 MODELING PHOTO-SWITCHING BEHAVIOR

The true photo-switching behavior of the fluorophore is a continuous time stochastic phenomenon. However, an experimenter can only ever observe a discretized manifestation of this by imaging the fluorophore in a sequence of frames. These frames are regarded as a set of sequential exposures of the fluorophore and the observed discrete time signal indicates whether the fluorophore has been observed in a particular frame. It is the continuous time process on which we wish to draw inference based on the observed discrete-time process indicating whether the fluorophore was observed in a frame. In this section we first present the continuous time Markov model of the true (hidden) photo-switching behavior, and then derive the observed discrete time signal, together with key results on its statistical properties.

### 2.1 Continuous time

We model the true photo-switching effect of the fluorophore as a continuous time Markov process, 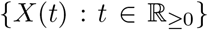 with discrete state space 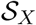. This is a stochastic process which satisfies the Markov property

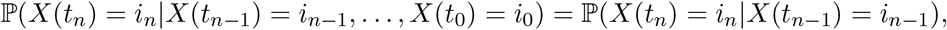

for any sequence of times *0* ≤ *t*_0_ < *t*_1_ < … < *t_n_* < ∞ and any sequence of states 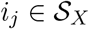 for *j* = 0, …, *n*.

In this paper we consider a general model for {*X*(*t*)} that can accommodate the numerous mechanisms of photo-switching utilized in standard SMLM approaches such as (F)PALM and (d) STORM. Specifically, this model consists of a photon emitting (On) state 1, *m*+1 non photon emitting (dark/temporary off) states 0_0_, 0_1_, …, 0_*m*_, where 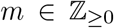, and a photobleached (absorbing/permanently off) state 2. In order to accommodate for the *m* = 0 case when we have a single dark state, we use the notational convention that state 0_0_ ≡ 0. The model, illustrated in Figure 4, allows for transitions from state 1 to the multiple dark states (from a photochemical perspective, these can include triplet, redox and quenched states). These dark states are typically accessed via the first dark state 0 (reached as a result of inter-system crossing of the excited S_1_ electron to the triplet T_1_ state - see Figure 2a). Further dark states 0_*i*+1_, *i* = 0, …, *m* − 1, are accessible by previous dark states 0_*i*_ (by, for example, the successive additions of electrons forming radical anions (Van de Linde et al., 2010)). We allow the On state 1 to be accessible by any dark state and we consider the most general model in which the absorption state 2 is accessible from any combination of other states (Vogelsang et al., 2010; Van de Linde and Sauer, 2014; Ha and Tinnefeld, 2012).

**Figure 4:**
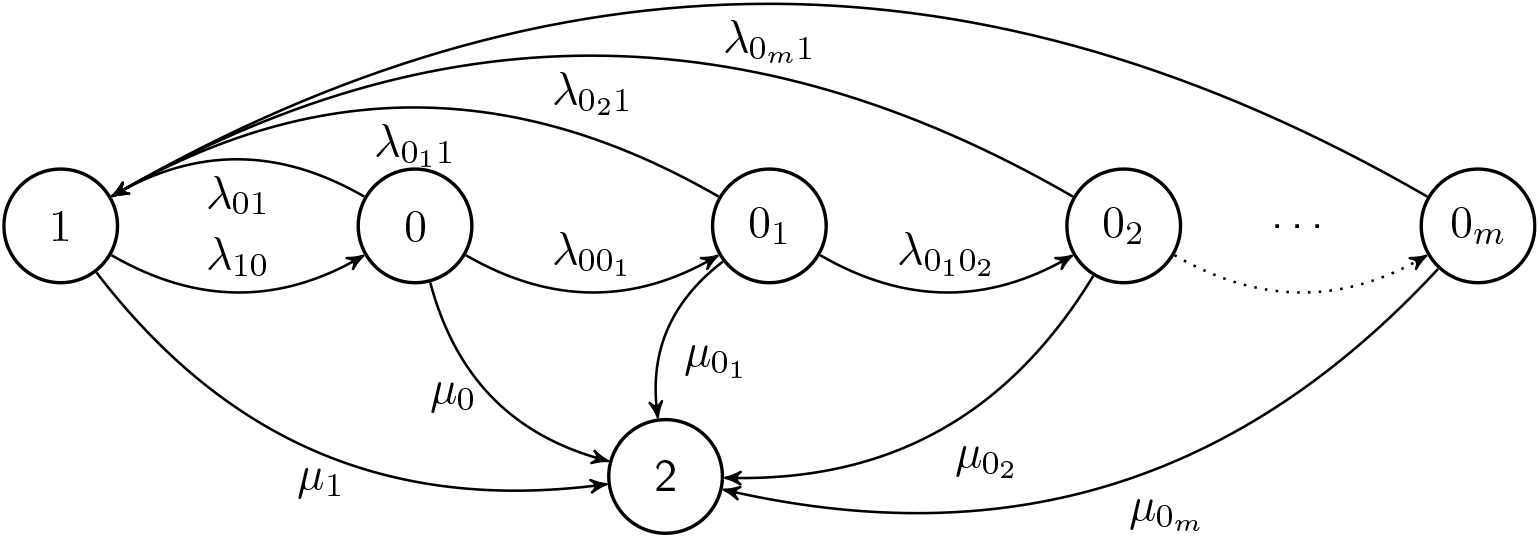
General *m* + 3 state 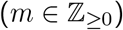 model of a fluorophore.

The state space of {*X*(*t*)} is 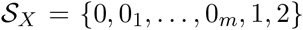 and is of cardinality *m* + 3. We denote *λ_ij_* to be the transition rate between states *i* and *j* and *μ_i_* to be the absorbing rate from state *i* to 2, where *i*, 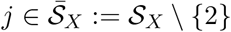.

The generator matrix for {*X*(*t*)} is therefore given as

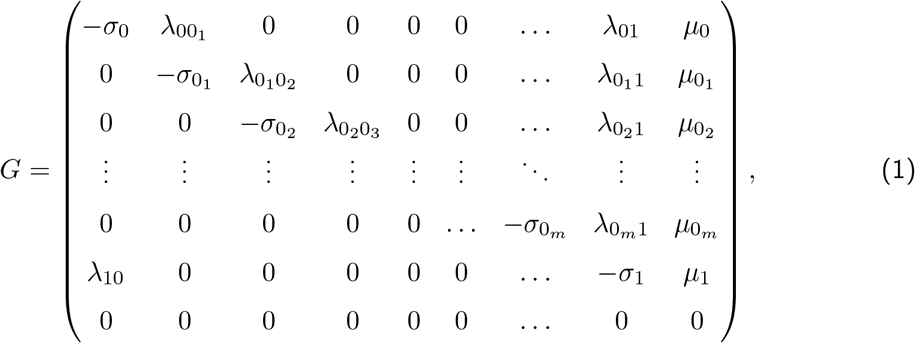

where *σ*_0_*m*__ = *λ*_0_*m*_1_ + *μ*_0_*m*__, *σ*_1_ = *λ*_10_ + *μ*_1_ and when *m* > 0, *σ*_0_*i*__ = *λ*_0_i_0_*i*+1__ + *λ*_0_*i*_1_ + *μ*_0_*i*__, for *i* = 0, …, *m* − 1. For full characterization, we define its initial probability mass ***ν***_*X*_ := (*ν*_0_ *ν*_0_1__ … *ν*_0_*m*__ *ν*_1_ *ν*_2_)^⊤^ with 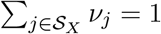. The most commonly occurring in practice is with *ν*_1_ = 1. Moreover, although the case when 1 > *ν*_2_ > 0 may give rise to fluorophores that are never observed, for inference purposes, we discard all traces containing no observations (1s) of fluorophores and set *ν*_2_ = 0.

In this paper, we will refer to specific models (from that shown in Figure 4) in the form 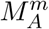 Here, *m* is as previously defined (denoting the number of multiple dark states beyond the 0_0_ state that are present in all models) and 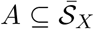 denotes the set of states from which the absorption state 2 is accessible. In particular, considering the three classical models presented in Figure 2: model (a) is 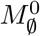: the *m* = 0 case where *μ*_0_ = *μ*_1_ = 0, model (b) is 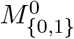: the *m* = 0 case where *μ*_0_, *μ*_1_ > 0, and model (c) is 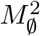: the *m* = 2 case where *μ*_0_ = *μ*_0_1__ = μ_1_ = 0.

### 2.2 Discrete time observation process

Having presented the continuous time model for the true photo-switching behavior, we will now introduce the model for the observed discrete time process and show how the transition rates given in (1) are not amenable to direct estimation.

The imaging procedure requires taking a series of successive frames. Frame *n* is formed by taking an exposure over the time interval [*n*Δ, (*n* + 1)Δ), where 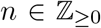. The constant Δ corresponds to the exposure time for a single frame, also known as the frame length. We define the *discrete* time observed process 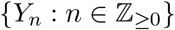, with state space 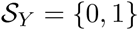, as *Y_n_* = 1 if the fluorophore (characterized by {*X*(*t*)}) is observed in frame *n* and equal to 0 otherwise. For the fluorophore to be observed in the time interval [*n*Δ, (*n* + 1)Δ) it must be in the On state 1 for a minimum time of *δ* ∈ [0, Δ). The value of *δ* is unknown and is a result of background noise and the imaging system’s limited sensitivity. We note that if {*X*(*t*)} exhibits multiple jumps to state 1 within a frame, then a sufficient condition for observing the fluorophore is that the total time spent in the On state exceeds *δ*. The *δ* = 0 case is the idealistic scenario of a noiseless system and perfect sensitivity such that the fluorophore is detected if it enters the On state for any non-zero amount of time during the exposure time Δ.

We formally define the observed process as

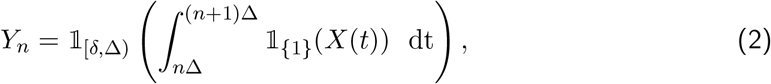

where 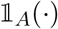 is the indicator function such that 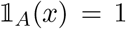 if *x* ∈ *A* and is zero otherwise. Figure 5 illustrates the manifestation of the discrete time signal {*Y_n_*} from the continuous time signal {*X*(*t*)}.

**Figure 5:**
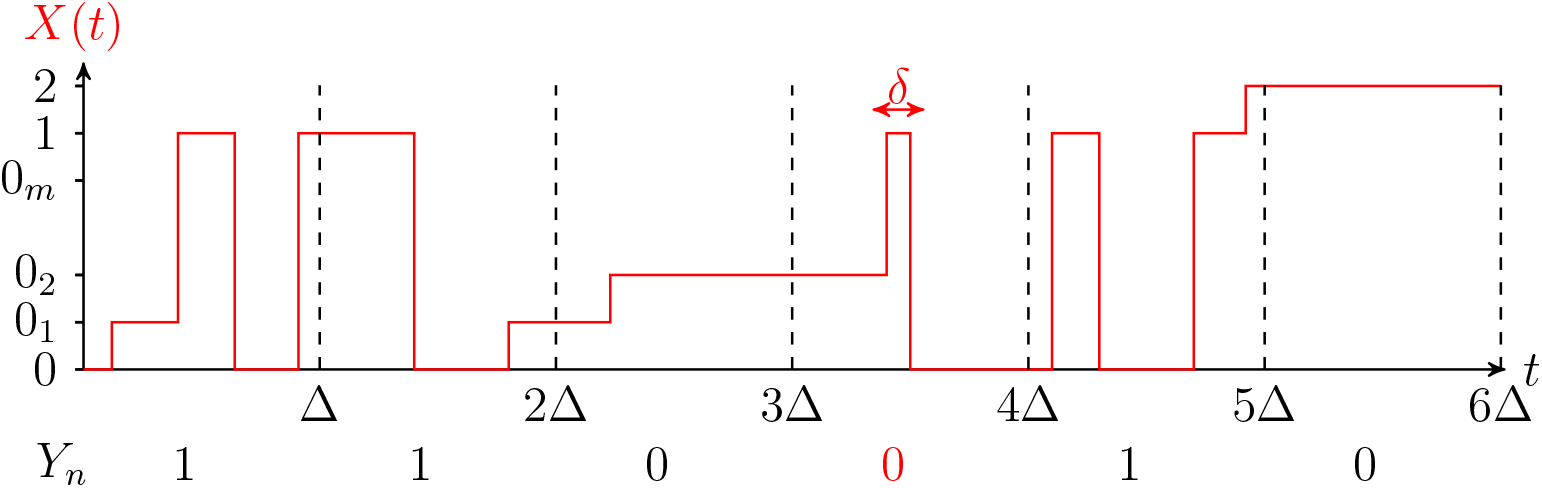
Illustration of how the states for *Y_n_* derive from the process *X*(*t*)

### 2.3 The inference problem

The inference problem is two-fold. Firstly for a given model, the aim is to estimate the unknown parameters

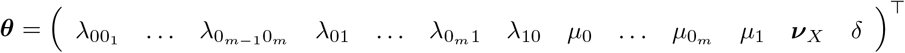

from a finite length realization of {*Y_n_*}. Crucially, it is shown in the Supplementary Materials Section S1 that {*Y_n_*} does not exhibit the Markov property (of any order) for any 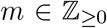, and for any Δ and *δ* such that Δ > *δ* ≥ 0. The non-Markovianity excludes classical inference methods and motivates the use of a Hidden Markov Model (HMM), with a likelihood based approach for estimating ***θ***.

Beyond this, it may be the case that the true model (characterized by its number of dark states) is unknown and may need to be selected in addition to estimating the unknown parameters. We tackle both of these problems in the next section.

## 3 CHARACTERIZING PHOTO-SWITCHING BEHAVIOR

Hidden Markov models, first presented in Baum and Petrie (1966), relate a sequence of observations to the states of an unobserved or *hidden* Markov chain. The aim of building a hidden Markov model (HMM) is to allow inference on the hidden process using these observations. In its simplest form, an HMM assumes the propagation of both state and observed sequences to be in discrete time, and a general first order HMM assumes that the observation process 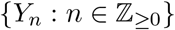 is related to a hidden first order Markov Chain 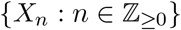 via an emission probability distribution 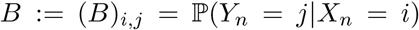, considered to be fully independent of the static parameters that characterize the probability distribution of state transitions 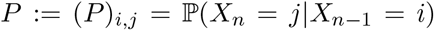. In this setting we say *B* and *P* are *decoupled*. For a sequence *y*_0_, *y*_1_, …, *y_N_* of observations from this model, the Baum-Welch re-estimation algorithm (Baum and Petrie, 1966; Baum and Eagon, 1967; Baum and Sell, 1968; Baum et al., 1970) is an EM type method that utilizes the forward-backward algorithm (see Levinson et al. (1983) for details) to optimize the likelihood function and compute maximum likelihood estimates of ***ν**_X_* (the probability mass of *X*_0_), *B* and *P*. This in turn can be used to estimate parameters of the emission and state transition probabilities. When the hidden Markov process and/or the observation process are of higher order, the HMM can be transformed to a general first order process (Du Preez, 1998; Lee and Lee, 2006; Ching et al., 2003) and Baum-Welch can be applied in the usual way. Readers are directed to MacDonald and Zucchini (1997) for a comprehensive review.

Whilst standard, first (or higher) order HMMs have been extensively studied and are most frequently used in applications, the rigid framework of being in discrete time with emission probabilities decoupled from state transition probabilities is not always suitable, as we will now show is the case for images formed by exposures over a time interval. We take time to carefully formulate the HMM suitable for this application, presenting what we call *transmission* (transmission-emission) matrices to capture the dependencies in the model. We then go on to provide a novel adaptation of the forward-backward algorithm to estimate ***θ***, the unknown parameters of our HMM in the case of a known state-space 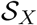 of the hidden process. We will then show how the Bayesian information criterion (BIC) can be used for model selection and parameter estimation in the case of an unknown state-space.

### 3.1 Photo-switching hidden Markov model

In this section we build an HMM for our observation process {*Y_n_*}, which we call the Photoswitching hidden Markov model (PSHMM). The first immediate reason as to why the standard set-up outlined above is inappropriate for this application is because the hidden Markov process {*X*(*t*)} evolves in continuous time. To deal with this, we need to adopt a time-discretization scheme for the hidden process. Analogously to Liu et al. (2015), we state that {*X*(*t*)} propagates in Δ-separated discrete time steps according to the transition probability matrix *P*_Δ_ = e^GΔ^, where *G* is given in (1). Our hidden process is therefore now represented by the discrete time Markov chain 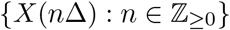.

When *Y_n_* depends solely on *X*(*n*Δ) (see Figure 6a) and the corresponding emission matrix *B* is decoupled from *P*, a continuous time EM algorithm (Liu et al., 2015) analogous to the Baum-Welch can be used to estimate ***ν**_X_*, *B* and *P*. However, this will be inappropriate in our setting for two related reasons. Firstly, we have shown in Section 2, specifically equation (2), that exposing images over a non-zero length of time means *Y_n_* depends on the full path of {*X*(*t*)} within the interval [*n*Δ, (*n* + 1)Δ). To correctly deal with this it is necessary to construct the HMM to consider dependence between *Y_n_* and both *X*(*n*Δ) and *X*((*n* + 1)Δ) (see Figure 6b). Secondly, this construction of {*Y_n_*} in (2) means the emission probabilities are clearly dependent on the static parameters ***θ*** of the hidden process and are therefore coupled with *P*. The EM procedures highlighted above require decoupled *B* and *P* so that at each step the quasi-likelihood can be optimized separately. To the best of our knowledge, methods for dealing with coupled systems have not been dealt with in the literature. While an EM algorithm could be used for a coupled system, analytic forms for the update steps would in general be intractable, leading to numerical maximization procedures at each iteration, thereby increasing computational complexity. We will now formally characterize the PSHMM and provide a novel method for estimating the unknown static parameters in the case of a coupled system.

**Figure 6:**
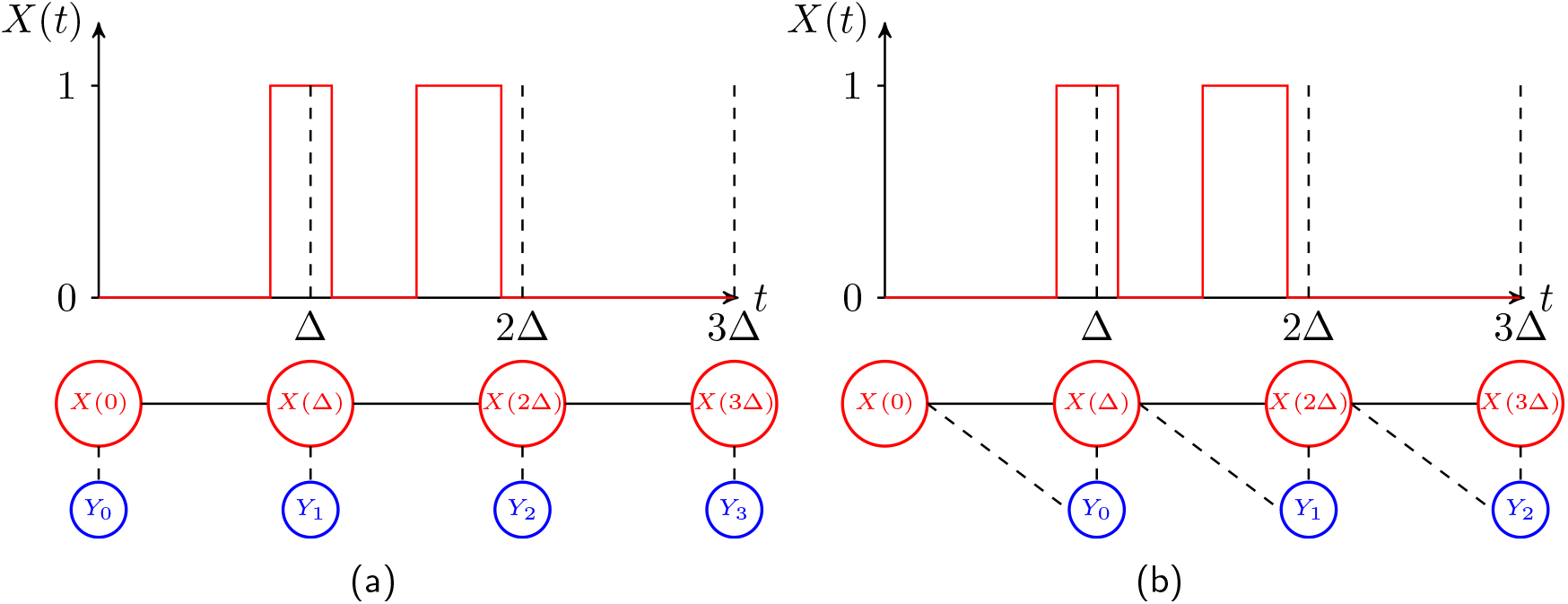
Illustration of the HMM setup. (a): Traditional HMM where observed state is dependent on current hidden. (b): Our HMM where observed state depends on both the current and past hidden states.

#### 3.1.1 Formal characterization of the PSHMM

Formally, we characterize our PSHMM with

1. an initial probability vector ***ν**_X_* = (*ν*_0_ *ν*_0_1__ … *ν*_0_*m*__ *ν*_1_ *ν*_2_)^⊤^ where 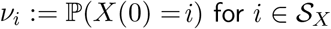;
2. Transmission matrices

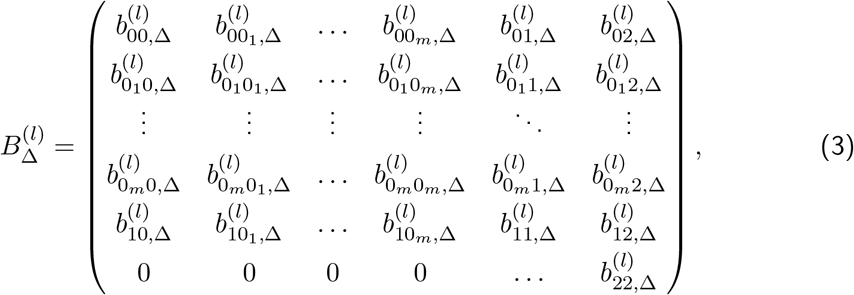

where

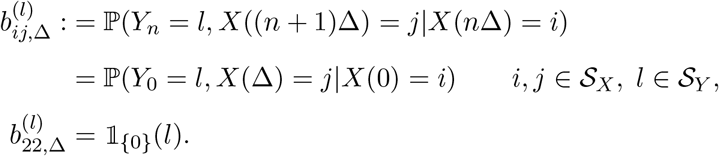

These transmission matrices combine the transition and emission probabilities, thereby allowing us to account for a coupled system. The full mathematical formulation for deriving their forms involves conditioning on the number of jumps from all *m* + 1 dark states within the interval [0, Δ). From this, we use Laplace transforms and the distributions of state holding times to iteratively compute matrices that converge to our set of transmission matrices. A more detailed explanation of this methodology, along with full derivations and expressions is presented in Supplementary Materials Section S2. Furthermore, an algorithm (Algorithm 1) detailing all computational steps to evaluate these matrices suitable for any 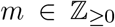 (any number of multiple dark states) can be found in Appendix A.

### 3.2 Estimating unknown parameters of the PSHMM

We now provide an algorithm for estimating the unknown parameters ***θ*** of the PSHMM, which utilizes a suitable adaptation of the forward-backward dynamic programming algorithm (Rabiner (1989)), making use of the transmission matrices in (3).

Let ***y*** = (*y*_0_ *y*_1_ … *y_N_F−1__*)^⊤^ be the sequence of observations across *N_F_* frames for a single photo-switching fluorophore. We define the forward-backward probabilities as

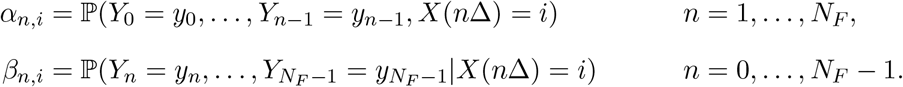

For each such *n*, we define the forward-backward vectors ***α**_n_* = (*α*_*n*,0_ … *α*_*n*,0_*m*__ *α*_*n*,1_ *α*_*n*,2_)^⊤^ and ***β**_n_* = (*β*_*n*,0_ … *β*_*n*,0_*m*__ *β*_*n*,1_ *β*_*n*,2_)^⊤^ Using this notation, we can show that 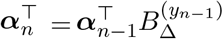 for *n* = 2, …, *N_F_* and 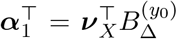 when *n* = 1. This yields the following recursion formula

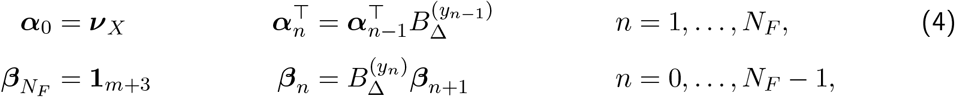

where **1**_*m*+3_ is the (*m* + 3) × 1 vector of ones. It now follows that the likelihood of observation vector ***y*** given parameter vector ***θ*** is 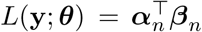 for all *n* = 0, …, *N_F_*. In particular, we have 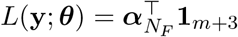, which can be readily computed using the transmission matrices together with recursive computation for 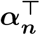 as indicated in (4). In the situation where we have *N_E_* ≥ 1 independent photo-switching fluorophores, the log-likelihood is given by

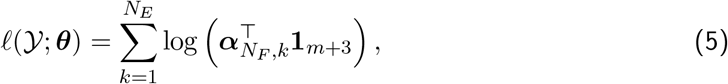

where 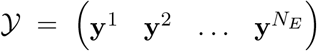 and ***α**_N_F_,k_* is the forward probability vector for emitter *k* = 1, …, *N_E_*. Maximizing (5) with respect to ***θ*** can be done either through numerically approximating derivatives or by using derivative-free optimization, for example with the Nelder-Mead algorithm. A discussion on multimodality and choosing a starting point for optimization can be found in the Supplementary Materials Section S3.

#### 3.2.1 Accounting for false positive observations

Occasionally, random peaks in the background noise may exceed the threshold value used to determine a fluorophore in the On state, resulting in a false positive identification of the fluorophore. For experiments conducted over a large enough number of frames, this false positive rate may become significant in the observed process {*Y_n_*}.

Specifically if *α* ∈ [0, 1] denotes the probability of falsely observing a fluorophore, assumed independent of the general observation process, then we may use the updated transmission matrices

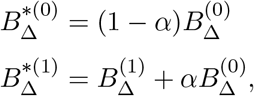

in the evaluation of the log-likelihood 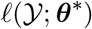 in (5). This would thus involve estimating ***θ***\* = [***θ***^⊤^ *α*]^⊤^ from the observations 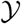.

### 3.3 Bootstrapping scheme

When only one experiment is conducted to produce an *N_F_* × *N_E_* dataset 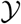, a single prediction 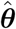 is obtained. In this circumstance, a bootstrapping scheme can be used to gain approximate confidence intervals for ***θ***.

In the same manner as is presented in Efron and Tibshirani (1993), we generate *R* (typically large) bootstrap datasets 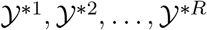 each consisting of re-sampled (with replacement) columns of 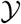. From each dataset, we acquire bootstrap replicated parameter estimates 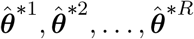 using the same PSHMM maximum likelihood procedure used to obtain 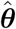. For 0.5 < *p* < 1, letting 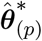 and 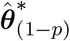 be the 100 · *p*th and 100 · (1 − *p*)th empirical percentiles from 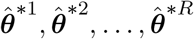 (with each component being the percentile from its corresponding parameter), a percentile bootstrap interval of length 1 − 2*p* is given by (see Efron and Tibshirani (1993))

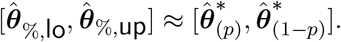

### 3.4 Model selection

In order to determine the *unknown* number of multiple *m* dark states, we may use the Bayesian information criterion (BIC) to determine the most likely model given data 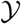. This particular model selection procedure is chosen due to the incorporation of the log-likelihood function which can be readily and easily computed, and further due to its penalization of over-fitting.

The BIC is defined in our context as being 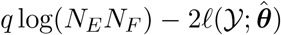, where *q* denotes the number of unknown parameters estimated in ***θ*** and 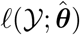 denotes the maximized log-likelihood using the maximum likelihood estimates 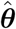. This criterion can be computed among all suitable models, with the most preferred being chosen as that with the smallest BIC value.

## 4 SIMULATION STUDY

In order to test the performance of parameter estimation against ground truth, synthetic imaging data of photo-switching fluorophores was simulated. We being our focus on the model 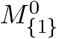, since for many practical applications the life-times of further dark (in particular the triplet (*T*_1_)) states is short relative to Δ. As such, this dark state has been considered as part of the meta-stable On state (Ha and Tinnefeld, 2012; Vogelsang et al., 2010). Since the predominant pathway to absorption is via the triplet state, this allows for a simplified model to be used in which the absorption state 2 is only accessible from state 1. Given the popularity of this model and its ease of analysis, we have derived the exact solution of the corresponding transmission matrices (see Supplementary Materials Section S2).

Details on the image simulation method and how the discretized state sequences were extracted can be found in the Supplementary Materials Section S4. Global parameter values are also noted. The extracted state sequences were analyzed using an implementation of Algorithm 1 (see Appendix A for theoretical details). The resulting parameter estimates were compared to estimates derived from the exponential fitting method, which was extended in this study to allow the calculation of absorption rates (see Supplementary Materials Section S3).

Table 2 (see Appendix B) shows estimated parameter statistics over 16 image simulation studies with 100 replicates (datasets) per study. Rate parameters ***θ***, were chosen to cover a range of observed behaviors of organic fluorophores and fluorescent proteins (Dempsey et al., 2011) with *N_E_* = 100 fluorophores per study. The number of frames *N_F_* in each study was adjusted to normalize against the average number of transitions predicted from ***θ***. Scatterplots of these rate estimates are presented in Figure 7. It is evident that the PSHHM yields estimates with much lower bias and root mean squared errors (RMSE) when compared to the exponential fitting method, although they have a tendency to increase as transition and absorption rates are increased. The reported 95% simulated intervals contain true parameter values across all studies from the PSHMM estimates and further highlight the bias in estimates obtained from the exponential fitting.

For experimenters, the effect of imaging parameters on the performance of the estimators is of particular interest and importance. Further simulation studies carried out under model 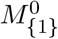 highlight the consistency in both accuracy and precision of the PSHMM estimator across a range of different experimental conditions. This is in particular emphasized in Figure 8, where we compare PSHMM with exponential fitting rate estimates when we vary the emission intensity of the fluorophores (measured in the mean number of photons each emits when in the On state for time Δ). Further investigation of other parameters, including the frame length (Δ), the number of frames (*N_F_*) and the detection threshold (proportional to *δ*) under this model, are provided in the Supplementary Materials Section S4. It is noted that across the full range of relevant parameters tested, the PSHMM estimator performs significantly better than exponential fitting.

**Figure 7:**
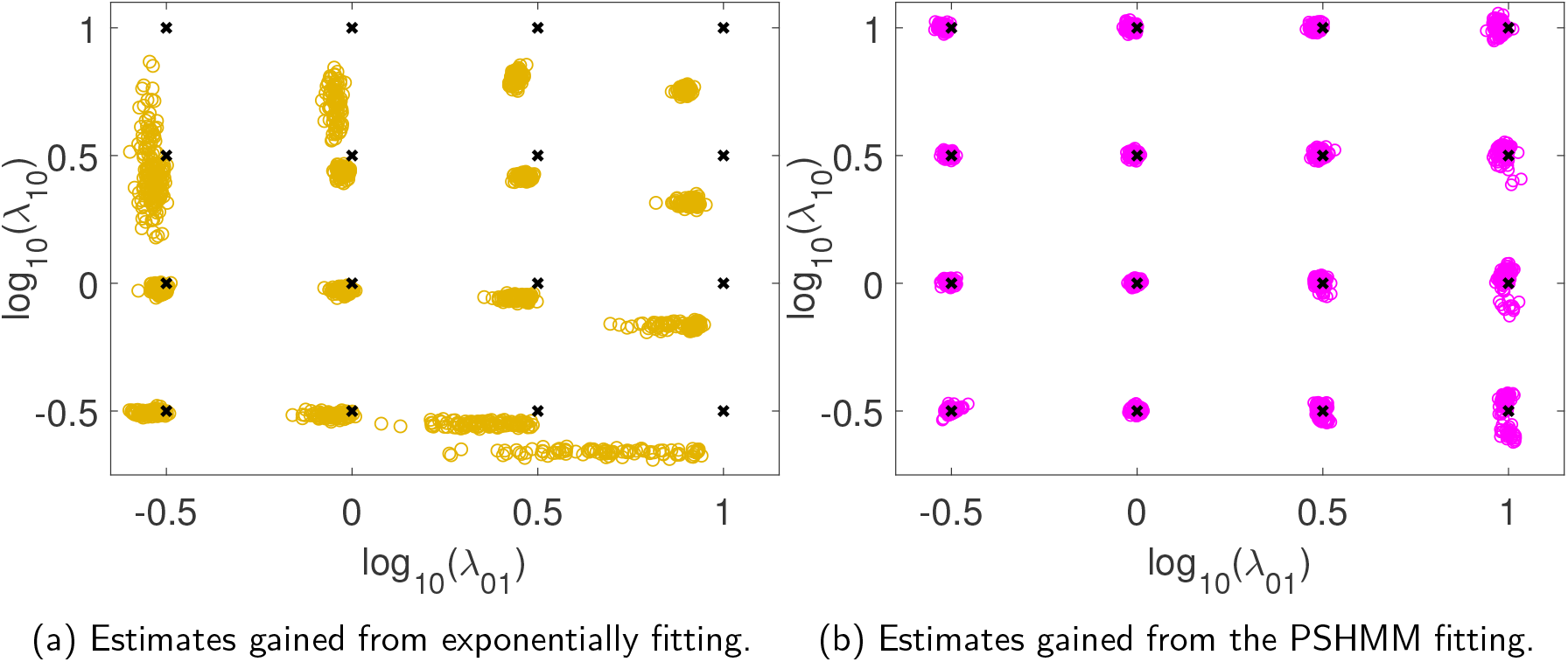
Estimates of log_10_(*λ*_01_) and log_10_(*λ*_10_) simulated from model 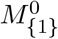 using both exponential fitting (7a) and PSHMM fitting (7b) are plotted in dark yellow and pink respectively. True rates are plotted as black crosses. Estimates for the absorption rate *μ*_1_, along with means, RMSEs and 95% simulated intervals are given in Table 2 (see Appendix B).

**Figure 8:**
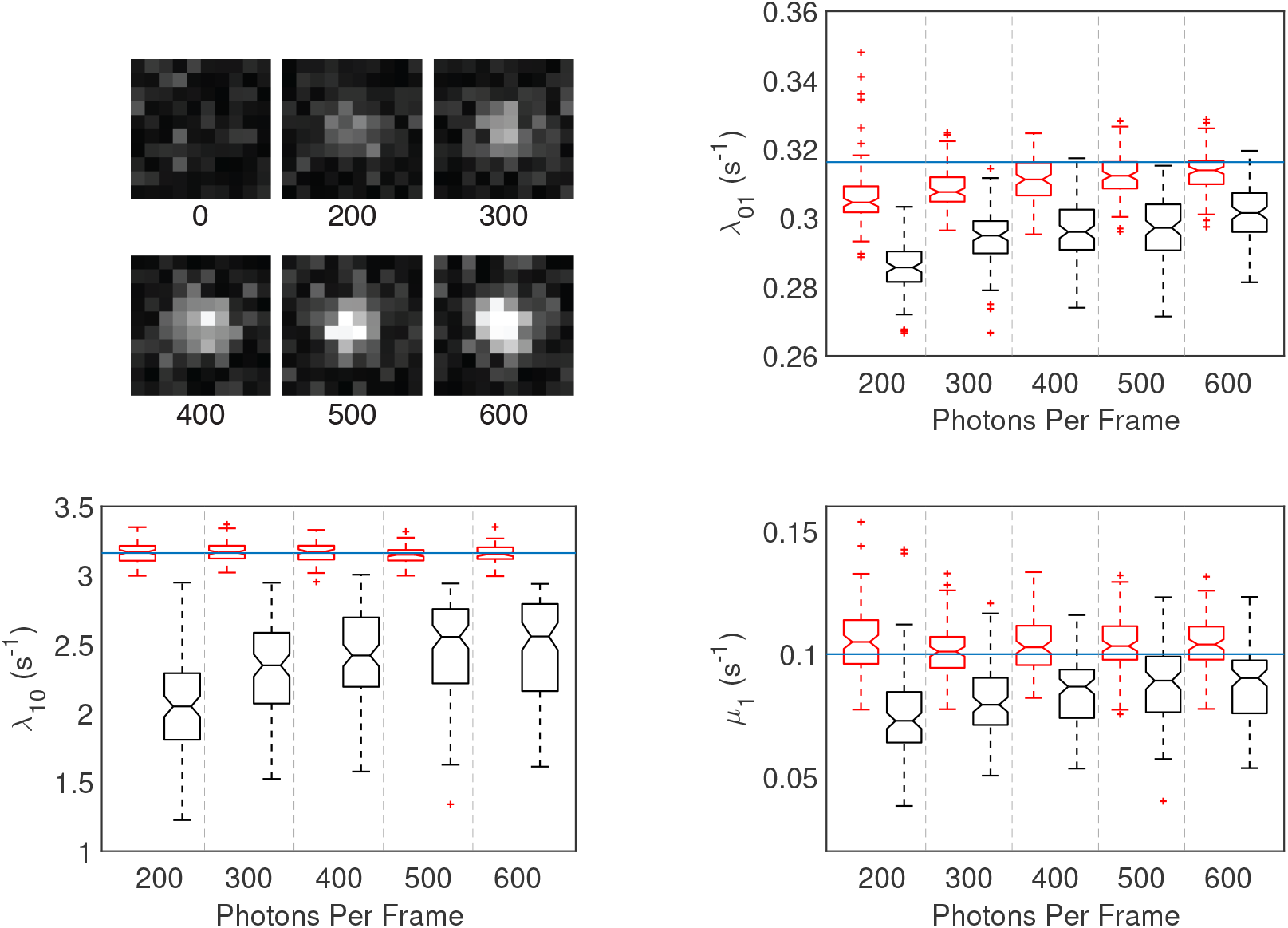
Top Left: Examples of single simulated frames at the indicated number of photons per frame (Supplementary Materials Section S4). Boxplots showing quantiles from estimates of *λ*_01_, *λ*_10_ and *μ*_1_ from both exponential fitting (black) and PSHMM fitting (red) are plotted against increasing photons per frame. *N_F_* = 9872 for all simulations. True rates given by the blue line.

In order to assess the accuracy of parameter estimates for the extended models *m* = 1 and *m* = 2 over fast, medium and slow switching scenarios, additional simulations were performed by directly sampling the continuous time processes {*X*(*t*)} and extracting the observation sequences 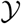 as in (2), using fixed values of ***θ***. Results from the analyses of these simulations are shown in Tables 3 and 4 in Appendix B. While it is evident that the estimates for *λ*_0_*m*_0_*m*+1__ and *λ*_0_*m*+1_1_ incur greater bias as m increases, the 95% simulated intervals predominantly cover true parameter values, albeit over a larger area due to the increase in the RMSEs. As is seen when *m* = 0, the exponential fitting method performs less well, yielding much higher bias and RMSEs for particular parameter values. The multimodality of the likelihood surface (see Supplementary Materials Section S3), especially in the case *m* = 2 invokes poorer estimates from both fitted methods.

Finally, using these simulated datasets, the BIC was used in model selection from the set of proposals 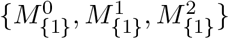 (i.e. under the assumption that the absorption state was known to only be accessible by the On state). Applying model selection to the 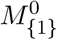 dataset used to estimate parameters in Table 2 results in the true state model being chosen in all (100%) cases. 100 datasets, each for *m* = 0,1,2 were generated for studies 2, 17 and 20 with 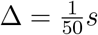 and *N_E_* = 300. These results presented in Table 1 demonstrate the accuracy of selecting the correct model.

**Table 1:** Confusion table showing the empirical percentage of models predicted from three candidates: 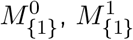 and 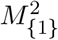 under simulation studies 16, 19 and 20 (see Tables 2, 3 and 4 in Appendix B), with *N_E_* = 300, 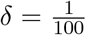 and 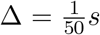. 100 datasets from each study were generated and the BIC used to select the best fitted model.

| Predicted →<br>True ↓ | $M_{\{1\}}^0$ | $M_{\{1\}}^1$ | $M_{\{1\}}^2$ |
| --- | --- | --- | --- |
| $M_{\{1\}}^0$ | 100 | 0 | 0 |
| $M_{\{1\}}^1$ | 0 | 98 | 2 |
| $M_{\{1\}}^2$ | 0 | 1 | 99 |

## 5 APPLICATION TO ALEXA FLUOR 647 DATA

In this section we apply the method presented in this paper to the data analyzed with the exponential fitting method in Lin et al. (2015). The details, including experimental methods, can be found in this reference. In summary, antibodies labeled with Alexa Fluor 647 at a ratio of 0.13−0.3 dye molecules per antibody were sparsely absorbed to a cover slip and imaged by Total Internal Fluorescence microscopy to investigate the effect of eight different laser intensities on the photo-switching behavior of Alexa Fluor 647. The study contains 27 experiments with differing combinations of laser intensity and frame rate. These values, together with the number of emitters detected and the number of frames over which they were imaged is summarized in Table 5 of Appendix B. For each photo-switchable molecule detected the discrete observation trace, indicating if the emitter was observed in each frame, was extracted (see Supplementary Materials Section S4). In all experiments, the true model and its associated parameters were unknown. Subsequently, we will show comparisons between estimates from both the PSHMM and modified exponential fitting methods ^1^.

Initially, the BIC model selection criterion as outlined in Section 3.4 was used to select the most suitable model for the data from the range of models 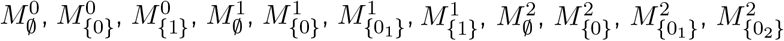 and 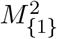, with the model 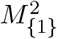 being selected on all (100%) occasions. This supports Lin et al. (2015), who hypothesize this, with bleaching model and assume the 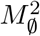 (without bleaching) model for rate estimates gained from exponential fitting. PSHMM maximum likelihood estimates were then computed for the unknown parameter vector ***θ***\* =(*λ*_00_1__ *λ*_01_ *λ*_0_1_0_2__ *λ*_0_1_1_ *λ*_0_2_1_ *λ*_10_ *μ*_1_ ***ν**_X_ δ α*)^⊤^ for each of the 27 datasets. Associated with these, 95% bootstrapped intervals were computed using the method in Section 3.3 (*R* = 100 due to computational intensity). The results are shown in Figures 9 and 10. Comparisons with exponential fitting bootstrapped re-estimates (where ***ν**_X_*, *δ* and *α* are not estimable in this setting) are also shown.

**Figure 9:**
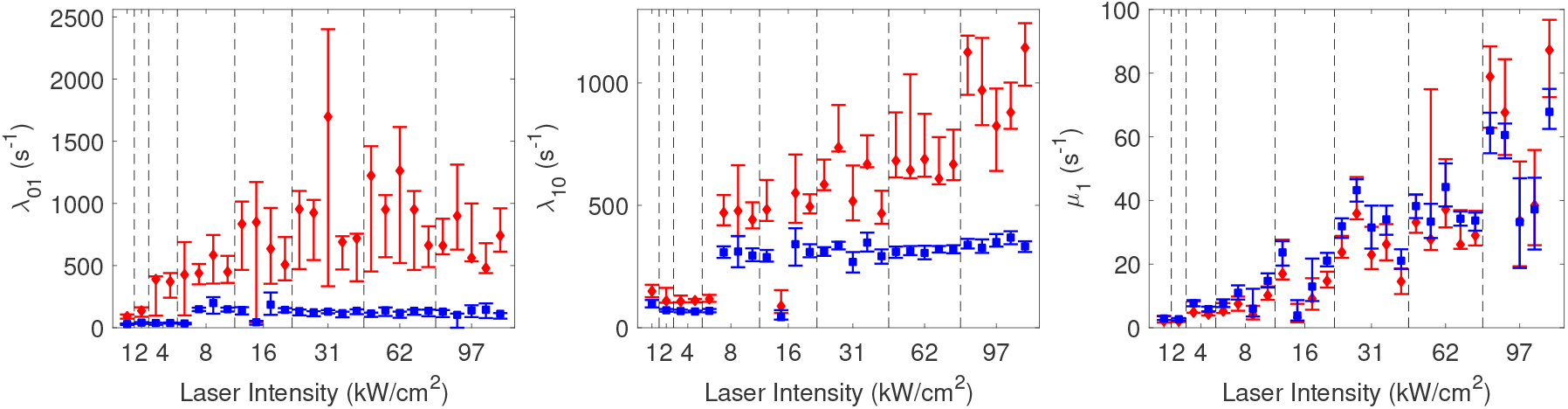
Rate predictions and associated 95% bootstrap confidence sets are shown for *λ*_01_ (left), and *λ*_10_ (middle) and *μ*_1_ (right) against increasing laser intensity (see Table 5 in Appendix B for exact values). Intervals in blue correspond to those from exponential fitting and those in red correspond to those gained from the PSHMM. Point estimates from each of the 27 datasets are given by the diamond (PSHMM) or square (exponential).

The results indicate that the exponential fitting predicts a much slower switching scenario for the Alexa Fluor 647 antibodies, with many estimates shown to be several orders of magnitude below those predicted by the PSHMM. This resembles the conclusions reached from the results of the simulation studies as described in Section 4 and are thought to occur as a result of the exponential fitting method missing events within frames. Incidentally, the higher variance of predictions from both methods are shown to be reported at higher laser intensities, where faster switching of fluorophores is promoted. This is especially pronounced in some particularly large simulated confidence sets for the exponential fitting estimates of *λ*_0_1_0_2__ and *λ*_0_2_1_ (see Figure 10).

**Figure 10:**
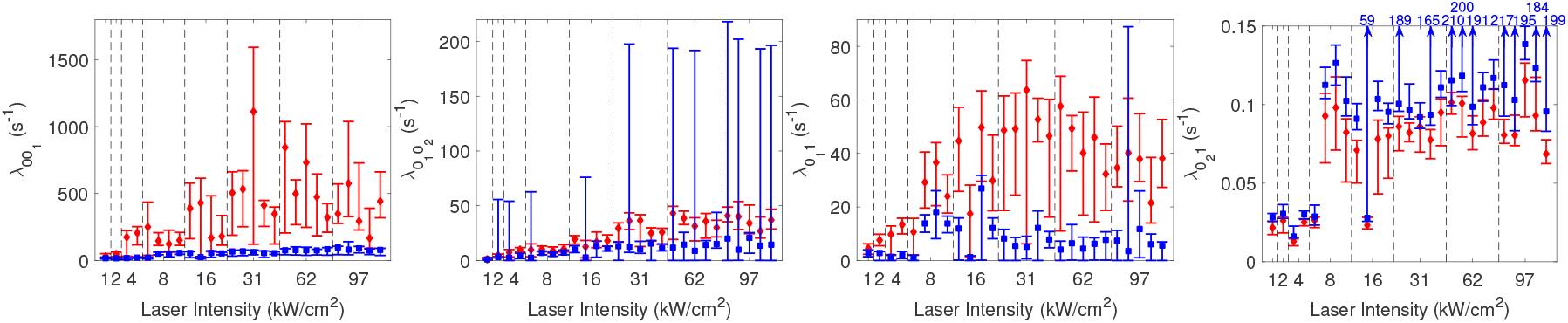
Rate predictions and associated 95% bootstrap confidence sets are shown for rates (in order from left to right) *λ*_0_1_1_, *λ*_0_1_0_2__, *λ*_0_1_1_ and *λ*_0_2_1_, plotted against increasing laser intensity (see Table 5 in Appendix B for exact values). Intervals in blue correspond to those from exponential fitting and those in red correspond to those gained from the PSHMM. Point estimates from each of the 27 datasets are given by the diamond (PSHMM) or square (exponential).

## 6 SUMMARY AND DISCUSSION

Accurate measurement of fluorophore photo-switching rates has the potential to enable tailored design of single molecule localization microscopy experiments to specific requirements. For example, one may wish to select a fluorophore and photo-switching environment to achieve the rapid photo-switching at low laser intensities required for live-cell samples. Alternatively, one may wish to promote long off times required for densely packed samples. Furthermore, precise estimates of photo-switching rates has the potential to advance data processing methods used in single molecule localization microscopy imaging, enabling more accurate image reconstruction and aiding proper quantitative analysis. For this purpose, we have presented a method for characterizing the photo-switching kinetics of fluorophores from a sequence of images.

For the most general continuous time photo-switching model, we have carefully defined the observation process and linked it to the hidden continuous time photo-switching behavior that we wish to infer upon. From this, we have formulated a hidden Markov model to link the observations to the continuous time photo-switching model. Importantly, images being formed by exposing the camera over a non-zero time interval violates the traditional assumption placed on HMMs that the emission and transition probabilities are decoupled. To tackle this, we have introduced transmission matrices that capture all the dependencies present in the model and provided a detailed scheme for computing them for any continuous time photo-switching model. A modification of the forward-backward algorithm tailored for these coupled HMMs has been presented and numerical maximization of the computed likelihood was performed to generate accurate estimates of the true photo-switching rates. Through a detailed simulation study, these were compared to estimates from an existing exponential fitting method. We found that our proposed method of parameter estimation is highly robust to a range of simulated experimental parameters including low signal-to-noise ratios and fast frame rates, frequently outperforming estimates from exponential fitting. Finally, we found that by using the BIC, it is possible to perform accurate model selection from a range of model proposals, thus providing a powerful new tool for chemists wishing to infer the number of quantum states a particular fluorophore can exist in. The model selection and estimation method presented in this paper was then applied to real data collected from the study of Lin et al. (2015). We provide strong evidence of a relationship between laser intensity and photo-switching rates and support the hypothesis that Alexa Fluor 647 has three off-states in addition to a photo-bleached state.

While this paper focuses on single molecule localization microscopy, the type of kinetic models discussed in this paper are unlikely to be unique to photo-switching fluorophores and super-resolution applications. Certainly, stochastic processes in which the observed signal depends on both the current and past states of a hidden process are likely to be a general feature of digital, discretized measurements of stochastic signals. This is particularly true in image processing where images are inevitably formed by exposing the camera’s sensor over a non-zero length time window. The coupling between the emission and transition probabilities of the HMM is a direct consequence of this exposure time, and therefore it is likely that the presented methodology for dealing with this will find use in imaging applications that are beyond the scope of this paper.

Further theoretical discussions and a comprehensive simulations and methods section, can be found in the Supplementary Materials.

## SUPPLEMENTARY MATERIALS

The supplementary materials supporting this paper contain detailed proofs and derivations regarding our method, discussions on its implementation and a section on further simulation studies, including exact details on the image analysis.

## CODE AND DATA

MATLAB code and imaging data sets (also attached separately) used for the algorithms presented in this paper, can be found at github.com/eakcohen/photoswitching.

## ACKNOWLEDGEMENTS

We would like to thank Prof Joerg Bewersdorf, Department of Cell Biology, Yale University for his help in making the Alexa Fluor 647 data available. This work was funded by grants from (R. Henriques) the Medical Research Council (MR/K015826/1), and Biotechnology and Biological Sciences Research Council (BB/M022374/1), and (R. Ober) National Institute of Health (R01 GM085575). N. Gustafsson funded by the Engineering and Physical Sciences Research Council (EP/L504889/1). L. Patel is funded by an Imperial College London President’s Scholarship.

## A APPENDIX: ALGORITHM TO COMPUTE TRANSMISSION MATRICES

Algorithm 1 presents the computation of transmission matrices 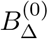 and 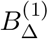 as defined in (3) (with full derivations found in the Supplementary Materials Section S2), suitable for any *m* ≥ 0. Here, we denote **0**_*n*_ and **1**_*n*_ to be the *n* × 1 vectors of zeros and ones respectively and ***I**_n_* to be the *n* × *n* identity matrix. We denote (*M*)_(*i*_1_:*i*_2_),(*j*_1_:*j*_2_)_ to be the matrix filled with rows *i*_1_ to *i*_2_ and columns *j*_1_ to *j*_2_ of any matrix *M*, and (*M*)_*i*_1_,*j*_1__ to be the (*i*_1_, *j*_1_)th entry of *M*. We use the ʘ notation to denote the Hadamard (element wise) product between two matrices.

### Algorithm 1 Compute transmission matrices 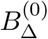 and 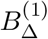

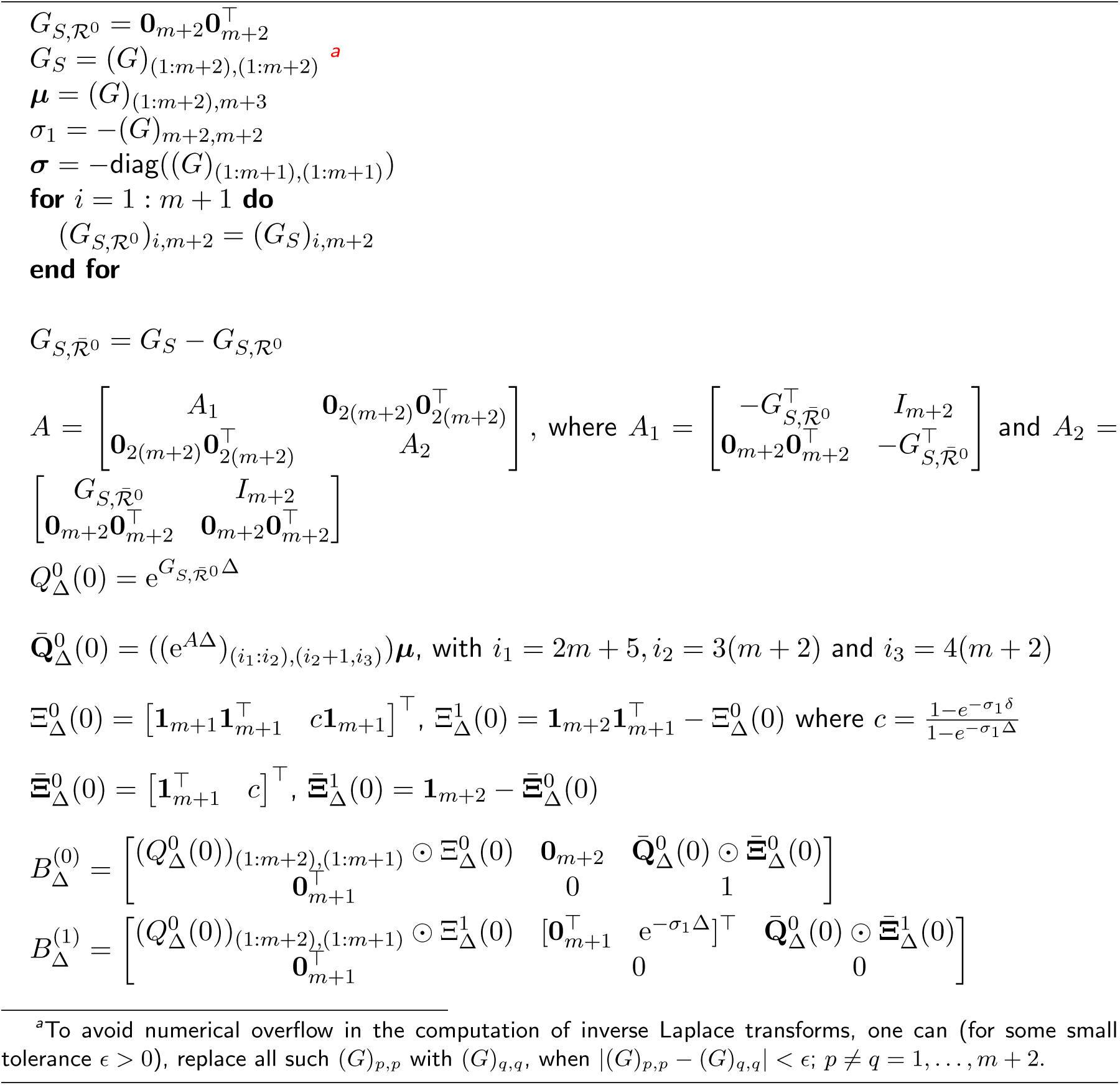
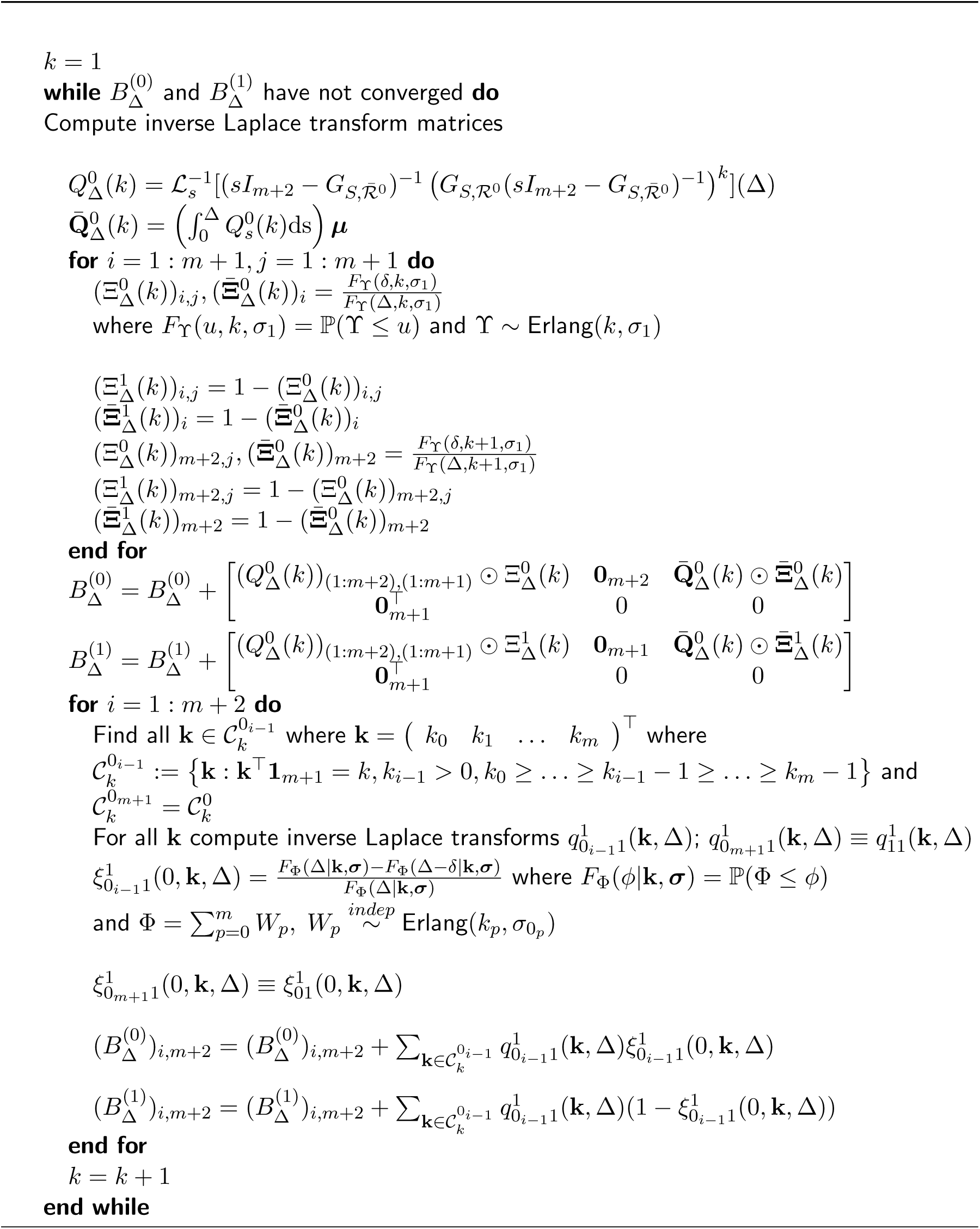

## B APPENDIX: RATE ESTIMATES

**Table 2:** Rate predictions of ***θ*** = (*λ*_01_ *λ*_10_ *μ*_1_)^⊤^ under model 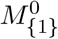 when 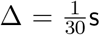, *δ*, *α* > 0 (unknown) and *N_E_* = 100 from both the PSHMM and exponential fitting (Exp) methods are presented along with means, root mean squared errors (RMSE) and 95% simulated intervals (S.I.). From both methods, log – log scatterplots of the photo-switching rates *λ*_01_ and *λ*_10_ are shown in Figure 7.

| Study | $N_F$ | $\theta$ | PSHMM<br>Mean | PSHMM<br>RMSE ($\times 10^{-2}$) | PSHMM<br>95% S.I. | Exp<br>Mean | Exp<br>RMSE ($\times 10^{-2}$) | Exp<br>95% S.I. |
| --- | --- | --- | --- | --- | --- | --- | --- | --- |
| 1 | 16800 | 0.3162 | 0.3192 | 0.9687 | (0.3045, 0.3412) | 0.2876 | 3.2931 | (0.2571, 0.3186) |
|  |  | 0.3162 | 0.3155 | 0.7613 | (0.3017, 0.3348) | 0.3094 | 0.8897 | (0.2987, 0.3222) |
|  |  | 0.0105 | 0.0110 | 0.2126 | (0.0080, 0.0173) | 0.0110 | 0.1051 | (0.0096, 0.0131) |
| 2 | 11151 | 0.3162 | 0.3152 | 0.6622 | (0.2986, 0.3271) | 0.3044 | 1.4688 | (0.2893, 0.3197) |
|  |  | 1 | 1.0026 | 1.9096 | (0.9602, 1.0403) | 0.9474 | 5.9482 | (0.8919, 0.9946) |
|  |  | 0.0333 | 0.0343 | 0.3470 | (0.0285, 0.0418) | 0.0347 | 0.3332 | (0.0300, 0.0428) |
| 3 | 9364 | 0.3162 | 0.3126 | 0.6751 | (0.3019, 0.3220) | 0.2994 | 1.9028 | (0.2791, 0.3157) |
|  |  | 3.1622 | 3.1638 | 6.5551 | (3.0491, 3.2826) | 2.4506 | 77.8492 | (1.7789, 2.9151) |
|  |  | 0.1054 | 0.1060 | 1.0206 | (0.0893, 0.1283) | 0.0889 | 2.1181 | (0.0627, 0.1173) |
| 4 | 8799 | 0.3162 | 0.3037 | 1.3963 | (0.2910, 0.3136) | 0.2840 | 3.3483 | (0.2669, 0.2997) |
|  |  | 10 | 9.9579 | 23.0254 | (9.5186, 10.4158) | 3.1910 | 690.8696 | (1.5220, 5.9107) |
|  |  | 0.3333 | 0.3471 | 3.7917 | (0.2934, 0.4186) | 0.1233 | 21.4912 | (0.0610, 0.2450) |
| 5 | 10962 | 1 | 0.9979 | 1.8580 | (0.9607, 1.0381) | 0.8963 | 12.4336 | (0.7390, 1.0060) |
|  |  | 0.3162 | 0.3163 | 0.7184 | (0.3017, 0.3317) | 0.3029 | 1.4713 | (0.2907, 0.3161) |
|  |  | 0.0105 | 0.0107 | 0.0993 | (0.0091, 0.0128) | 0.0112 | 0.1096 | (0.0097, 0.0129) |
| 6 | 5312 | 1 | 0.9962 | 1.8118 | (0.9602, 1.0311) | 0.9464 | 6.4432 | (0.8729, 1.0142) |
|  |  | 1 | 1.0006 | 1.7552 | (0.9623, 1.0355) | 0.9340 | 6.8768 | (0.8891, 0.9654) |
|  |  | 0.0333 | 0.0338 | 0.2904 | (0.0274, 0.0403) | 0.0347 | 0.2772 | (0.0296, 0.0403) |
| 7 | 3526 | 1 | 0.9850 | 2.3208 | (0.9492, 1.0165) | 0.9468 | 5.7823 | (0.9107, 0.9900) |
|  |  | 3.1622 | 3.1656 | 7.3336 | (3.0095, 3.3030) | 2.7115 | 46.1604 | (2.4988, 2.8888) |
|  |  | 0.1054 | 0.1062 | 1.0589 | (0.0839, 0.1274) | 0.1013 | 0.9895 | (0.0844, 0.1206) |
| 8 | 2961 | 1 | 0.9670 | 3.9061 | (0.9331, 1.0025) | 0.9053 | 9.7545 | (0.8524, 0.9451) |
|  |  | 10 | 9.9362 | 27.2081 | (9.4582, 10.4740) | 5.0187 | 504.3446 | (3.6585, 6.5223) |
|  |  | 0.3333 | 0.3500 | 3.9408 | (0.2808, 0.4209) | 0.2002 | 13.7743 | (0.1440, 0.2689) |
| 9 | 9116 | 3.1622 | 3.1546 | 6.8802 | (3.0410, 3.2910) | 2.3068 | 95.1931 | (1.6353, 3.0432) |
|  |  | 0.3162 | 0.3145 | 1.5272 | (0.2844, 0.3382) | 0.2793 | 3.7425 | (0.2694, 0.2901) |
|  |  | 0.0105 | 0.0107 | 0.1129 | (0.0089, 0.0133) | 0.0113 | 0.1253 | (0.0095, 0.0136) |
| 10 | 3466 | 3.1622 | 3.1275 | 7.4660 | (3.0121, 3.2771) | 2.8271 | 37.7579 | (2.4890, 3.0580) |
|  |  | 1 | 1.0042 | 4.0430 | (0.9027, 1.0673) | 0.8711 | 13.0043 | (0.8354, 0.9021) |
|  |  | 0.0333 | 0.0338 | 0.3699 | (0.0265, 0.0417) | 0.0350 | 0.3610 | (0.0287, 0.0415) |
| 11 | 1680 | 3.1622 | 3.1101 | 9.4936 | (2.9827, 3.3063) | 2.9176 | 25.6336 | (2.7468, 3.0751) |
|  |  | 3.1622 | 3.1774 | 9.1226 | (2.9934, 3.3673) | 2.5950 | 56.9611 | (2.4846, 2.6953) |
|  |  | 0.1054 | 0.1073 | 1.2117 | (0.0875, 0.1309) | 0.1050 | 0.9772 | (0.0892, 0.1254) |
| 12 | 1115 | 3.1622 | 3.0275 | 14.7255 | (2.9162, 3.1519) | 2.7851 | 38.1889 | (2.6624, 2.9165) |
|  |  | 10 | 9.9922 | 24.8620 | (9.5353, 10.4764) | 6.3522 | 365.9653 | (5.7218, 6.9261) |
|  |  | 0.3333 | 0.3485 | 3.9202 | (0.2874, 0.4364) | 0.2721 | 6.7863 | (0.2229, 0.3310) |
| 13 | 8532 | 10 | 9.9308 | 25.6393 | (9.4711, 10.4659) | 5.3340 | 506.7013 | (1.9767, 8.5991) |
|  |  | 0.3162 | 0.3163 | 4.4235 | (0.2421, 0.3674) | 0.2169 | 9.9483 | (0.2059, 0.2253) |
|  |  | 0.0105 | 0.0106 | 0.0893 | (0.0089, 0.0122) | 0.0111 | 0.0982 | (0.0096, 0.0125) |
| 14 | 2882 | 10 | 9.8578 | 29.5173 | (9.4399, 10.3920) | 7.7536 | 241.8453 | (5.5293, 8.7067) |
|  |  | 1 | 1.0264 | 10.5309 | (0.7836, 1.1608) | 0.6768 | 32.3653 | (0.6447, 0.7073) |
|  |  | 0.0333 | 0.0342 | 0.3583 | (0.0277, 0.0415) | 0.0355 | 0.3731 | (0.0299, 0.0421) |
| 15 | 1096 | 10 | 9.7339 | 38.3953 | (9.2222, 10.3385) | 8.1863 | 184.8409 | (7.3752, 8.6577) |
|  |  | 3.1622 | 3.2110 | 20.7343 | (2.4524, 3.4669) | 2.0539 | 110.9651 | (1.9708, 2.1564) |
|  |  | 0.1054 | 0.1095 | 1.2225 | (0.0897, 0.1335) | 0.1090 | 1.0413 | (0.0941, 0.1322) |
| 16 | 531 | 10 | 9.4992 | 55.7165 | (9.1001, 9.9554) | 7.9281 | 207.9020 | (7.5456, 8.2175) |
|  |  | 10 | 9.9053 | 54.4724 | (9.0157, 10.8772) | 5.6321 | 436.9608 | (5.4028, 5.8869) |
|  |  | 0.3333 | 0.3400 | 4.5056 | (0.2617, 0.4282) | 0.3044 | 4.0626 | (0.2602, 0.3643) |

**Table 3:** Rate predictions of ***θ*** = (*λ*_00_1__ *λ*_01_ *λ*_0_1_1_ *λ*_10_ *μ*_1_)^⊤^ under model 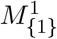 when 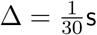, 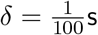, *α* = 0 and *N_E_* = 100 from both the PSHMM and exponential fitting (Exp) methods are shown along with means, root mean squared errors (RMSE) and 95% simulated intervals (S.I.).

| Study | $N_F$ | $\theta$ | PSHMM<br>Mean | PSHMM<br>RMSE ($\times 10^{-2}$) | PSHMM<br>95% S.I. | Exp<br>Mean | Exp<br>RMSE ($\times 10^{-2}$) | Exp<br>95% S.I. |
| --- | --- | --- | --- | --- | --- | --- | --- | --- |
| 17 | 11151 | 0.15 | 0.1519 | 1.6908 | (0.1192, 0.1945) | 0.1463 | 1.6566 | (0.1132, 0.1874) |
|  |  | 0.3 | 0.3006 | 0.8477 | (0.2837, 0.3178) | 0.2979 | 0.8373 | (0.2818, 0.3144) |
|  |  | 0.1 | 0.1004 | 0.4251 | (0.0928, 0.1090) | 0.1017 | 0.4528 | (0.0950, 0.1106) |
|  |  | 0.8 | 0.7995 | 1.2760 | (0.7750, 0.8240) | 0.7606 | 4.1190 | (0.7372, 0.7867) |
|  |  | 0.01 | 0.0100 | 0.1531 | (0.0073, 0.0136) | 0.0196 | 0.9666 | (0.0182, 0.0215) |
| 18 | 9364 | 0.35 | 0.3553 | 5.4361 | (0.2385, 0.4322) | 0.3278 | 5.3241 | (0.2401, 0.4294) |
|  |  | 1 | 1.0031 | 3.6825 | (0.9445, 1.0740) | 0.9514 | 5.8263 | (0.9002, 1.0113) |
|  |  | 0.3 | 0.2984 | 2.0116 | (0.2599, 0.3440) | 0.2918 | 2.1192 | (0.2535, 0.3342) |
|  |  | 2.3 | 2.2974 | 5.0064 | (2.2068, 2.3942) | 2.0382 | 26.4788 | (1.9520, 2.1115) |
|  |  | 0.1 | 0.1020 | 0.9824 | (0.0863, 0.1223) | 0.0950 | 1.0272 | (0.0808, 0.1140) |
| 19 | 7000 | 2 | 2.0333 | 18.1393 | (1.7516, 2.4518) | 2.1563 | 21.2523 | (1.8902, 2.4979) |
|  |  | 10 | 9.7821 | 54.4940 | (8.5534, 10.5310) | 6.9389 | 306.6856 | (6.5918, 7.3413) |
|  |  | 0.7 | 0.7109 | 4.8848 | (0.6364, 0.8341) | 0.6687 | 4.8472 | (0.6024, 0.7629) |
|  |  | 10 | 10.0021 | 63.6158 | (9.2232, 11.6534) | 4.9697 | 503.1383 | (4.7549, 5.1729) |
|  |  | 0.333 | 0.3380 | 7.2873 | (0.2035, 0.5619) | 0.2651 | 7.2969 | (0.2237, 0.3168) |

**Table 4:** Rate predictions of ***θ*** = (*λ*_00_1__ *λ*_01_ *λ*_0_1_0_2__ *λ*_0_1_1_ *λ*_0_2_1_ *λ*_10_ *μ*_1_)^⊤^ under model 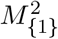 when 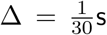, 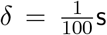, *α* = 0 and N_E_ = 100 from both the PSHMM and exponential fitting (Exp) methods are shown along with means, root mean squared errors (RMSE) and 95% simulated intervals (S.I.).

| Study | $N_F$ | $\theta$ | PSHMM<br>Mean | PSHMM<br>RMSE ($\times 10^{-2}$) | PSHMM<br>95% S.I. | Exp<br>Mean | Exp<br>RMSE ($\times 10^{-2}$) | Exp<br>95% S.I. |
| --- | --- | --- | --- | --- | --- | --- | --- | --- |
| 20 | 7000 | 2 | 2.0322 | 3.1417 | (1.7498, 2.3711) | 2.0535 | 2.1195 | (1.7933, 2.3121) |
|  |  | 10 | 9.8470 | 13.4172 | (9.2041, 10.4857) | 7.0420 | 878.8248 | (6.6792, 7.4758) |
|  |  | 0.2 | 0.2088 | 0.1033 | (0.1605, 0.2744) | 0.1762 | 0.1075 | (0.1289, 0.2128) |
|  |  | 0.7 | 0.6878 | 0.3748 | (0.5866, 0.8288) | 0.6635 | 0.3458 | (0.5703, 0.7536) |
|  |  | 0.01 | 0.0094 | 0.0005 | (0.0056, 0.0135) | 0.0149 | 0.0196 | (0.0128, 0.0176) |
|  |  | 10 | 9.6319 | 35.5731 | (8.7265, 10.5304) | 4.9131 | 2588.85 | (4.6687, 5.1636) |
|  |  | 0.333 | 0.3243 | 0.3224 | (0.2379, 0.4494) | 0.3206 | 0.1163 | (0.2616, 0.3844) |

**Table 5:** A description of the Alexa Fluor 647 datasets with reference to the laser intensities in kW/cm^2^ and frames sampled per second (or Δ^−1^) measured in s^−1^ used to characterize each of the 27 experiments. The *N_F_* × *N_E_* size of each dataset is also included.

| Dataset | Laser intensity | $\Delta^{-1}$ | $N_E$ | $N_F$ | Dataset | Laser intensity | $\Delta^{-1}$ | $N_E$ | $N_F$ | Dataset | Laser intensity | $\Delta^{-1}$ | $N_E$ | $N_F$ |
| --- | --- | --- | --- | --- | --- | --- | --- | --- | --- | --- | --- | --- | --- | --- |
| 1 | 1.0 | 200 | 275 | 49796 | 10 | 16 | 200 | 292 | 39703 | 19 | 62 | 800 | 443 | 29107 |
| 2 | 1.9 | 200 | 259 | 49533 | 11 | 16 | 800 | 305 | 29074 | 20 | 62 | 800 | 425 | 29551 |
| 3 | 3.9 | 200 | 335 | 49815 | 12 | 16 | 800 | 290 | 29145 | 21 | 62 | 800 | 425 | 29426 |
| 4 | 3.9 | 200 | 393 | 39758 | 13 | 31 | 800 | 617 | 29059 | 22 | 62 | 800 | 398 | 28989 |
| 5 | 7.8 | 200 | 340 | 39721 | 14 | 31 | 800 | 534 | 29778 | 23 | 97 | 800 | 454 | 29191 |
| 6 | 7.8 | 800 | 244 | 29418 | 15 | 31 | 800 | 515 | 29179 | 24 | 97 | 800 | 440 | 29198 |
| 7 | 7.8 | 800 | 230 | 29257 | 16 | 31 | 800 | 493 | 29400 | 25 | 97 | 800 | 436 | 29270 |
| 8 | 7.8 | 800 | 230 | 29438 | 17 | 31 | 800 | 456 | 29071 | 26 | 97 | 800 | 422 | 29295 |
| 9 | 16 | 800 | 437 | 29467 | 18 | 62 | 800 | 554 | 29327 | 27 | 97 | 800 | 414 | 29218 |

## Footnotes

1 We modified the exponential fitting algorithm used by Lin et al. (2015) to allow for the absorption parameter (see Supplementary Materials Section S3 for more details).

